# Automated Proofreading of Digitally Reconstructed Neural Morphology Enhances Accuracy, Scalability, and Standardization

**DOI:** 10.64898/2026.03.27.714818

**Authors:** Herve Emissah, Carolina Tecuatl, Giorgio A. Ascoli

## Abstract

**Background:** The rapid expansion of large-scale neuroscience datasets has increased the need for automated, accurate, and standardized quality control (QC). Manual proofreading of 3-dimensional neural morphology (SWC files) remains labor-intensive, error-prone, and non-scalable. We developed and evaluated a fully automated, machine-learning– driven QC pipeline to standardize neural reconstructions, detect and correct structural anomalies, and rectify dendritic labeling in pyramidal neurons.

**Methods:** We developed an end-to-end, cloud-deployed pipeline for automated QC, correction, and standardization of SWC-formatted neural morphologies. The framework integrates deterministic structural normalization, topology repair, geometric correction, quantitative morphometric analysis, and graph-based dendritic relabeling within a containerized React/Flask architecture deployed on Amazon Web Services. Rule-based algorithms systematically detect, classify, and correct structural irregularities including overlapping nodes, spurious side branches, non-positive radii, disconnected components, and anomalously long parent-child connections. A graph convolutional network, trained on Sholl-derived features from 20,500 pyramidal neurons, performs dendritic relabeling. Model training employed an 80/10/10 train–validation–test split with adaptive learning-rate scheduling and distributed execution across ten runs to evaluate stability and reproducibility. The pipeline generates images of the final product and computes quantitative morphometrics using L-Measure.

**Results:** All neuronal reconstructions were processed without manual intervention. Automated normalization and topology repair restored structurally coherent and biologically accurate morphologies suitable for quantitative analysis and visualization without data loss. Dendritic relabeling achieved a mean accuracy of 99.51%, consistent between validation and test sets, with class-weighted precision of 0.978, recall of 0.977, and F1-score of 0.977. Enforcing a single apical dendritic tree per neuron improved anatomical consistency without reducing classification performance. Distributed training completed all runs in approximately 25 hours, demonstrating scalability and reproducibility for large datasets.

**Conclusions:** We present a fully automated and cloud-scalable open-source pipeline for standardizing neural reconstructions and performing biologically consistent dendritic classification with near-perfect accuracy. The automated correction and relabeling procedures do not alter or compromise the size or unaffected morphological detail of the original SWC files, ensuring geometric fidelity and compatibility with downstream analysis tools. This open-access framework provides a robust foundation for high-throughput neural morphology curation and large-scale neuroanatomical analysis.

**Data Availability:** Processed results, model outputs, and source code are available at https://github.com/HerveEmissah/nmo_swc_standardization

The executable pipeline is accessible at https://swcstandardization.computational-neuromorpho.org

## 1. Introduction

Three-dimensional digital reconstructions of neuronal and glial morphology play a central role in modern neuroscience, enabling quantitative studies of neurite architecture, branching processes, network connectivity, and circuit organization across development, disease, and species [1-4]. Standardized neural morphology files, commonly stored in the SWC format, support large-scale neuroinformatics resources, computational simulations, and machine-learning-driven discovery pipelines [5-8]. The public repository NeuroMorpho.Org has now aggregated hundreds of thousands of neural reconstructions contributed by laboratories worldwide, reflecting rapid increases in the scale, resolution, and heterogeneity of morphological datasets [1,9]. As the volume of morphological data continues to grow, ensuring processing accuracy, consistency, and reproducibility has become a critical challenge for the neuroscience community [10-13].

Manual proofreading of neuronal reconstructions remains the predominant approach for resolving tracing artifacts, enforcing format validity, pruning anatomical errors, and labeling dendritic arbors such as apical and basal trees in pyramidal cells [14,15]. However, manual intervention is time-consuming, subject to inter-annotator variability, and increasingly impractical in the era of high-throughput imaging pipelines and large-scale connectomics initiatives [16-18]. Standardization tools exist to evaluate reconstruction quality and verify SWC compliance, yet these tools still require expert intervention, fragmented workflows, or non-scalable computing environments [5,6,10,19]. Likewise, automated structural checks do not fully address biological plausibility concerns such as anomalous branch geometry or dendritic mis-labeling, issues that directly impact downstream biological interpretation, morphometric analyses, and computational modeling [20-22].

Recent advances in machine learning, high-performance computing, and cloud-based ecosystems provide an opportunity to fundamentally modernize neural morphology quality control (QC). Graph-based learning frameworks, including graph convolutional neural networks (GCNs), have demonstrated strong performance in neuron classification and structural inference tasks by leveraging the topological nature of dendritic trees [23–26]. Yet, despite this technological progress, no end-to-end automated solution currently exists to standardize SWC files, correct structural irregularities, and apply biologically consistent dendritic labeling at scale within a unified, web-accessible, and user-friendly environment.

To address this gap, we designed, implemented, deployed, and evaluated a fully automated, cloud-based QC platform that integrates existing morphology-standardization utilities, custom correction algorithms, and a GCN-based dendritic relabeling model into a scalable cloud architecture. Our system enables researchers to upload neural reconstructions, automatically detect and correct structural anomalies, enforce SWC compliance, repair biologically implausible dendritic configurations, and assign apical and basal dendrite labels with high accuracy, all with minimal user intervention. The objective of this pipeline is to support large-scale neuroinformatics workflows while reducing manual burden and increasing reproducibility and standardization across laboratories and datasets.

## 2. Materials and Methods

We developed a web-based QC platform for standardized processing, structural correction, and automated dendritic relabeling of neural reconstructions encoded in SWC format and submitted by research groups worldwide to NeuroMorpho.Org. The platform integrates a React-based front end with a Flask backend, containerized using Docker and deployed on Amazon Web Services (AWS), enabling scalable and reproducible automated analysis workflows. Through the web-based application programming interface (API), users upload one or more SWC files, which are stored in a dedicated Source-Version directory within the containerized processing environment.

Upon upload, the backend verifies file presence, extension, structural validity, and admissible node types before initiating downstream processing. Invalid or malformed files are separated into quarantine folders, whereas valid reconstructions are assigned appropriate permissions and routed into the corresponding analysis pipelines. The system then executes three primary automated workflows: (i) structural standardization, (ii) long-connection detection and correction, and (iii) automated pyramidal dendritic relabeling. Each pipeline operates on the same SWC input but focuses on different aspects of morphological QC.

The first pipeline performs structural standardization to ensure consistent formatting and topology across uploaded reconstructions prior to further analysis. At the beginning of each processing run, temporary and output directories are cleared and recreated to provide a clean processing environment. Each SWC file is examined line by line to verify the expected seven-column format, valid parent-child relationships, and permitted node types. To maintain compatibility with normalization utilities and legacy datasets, certain branch-type labels are temporarily remapped during preprocessing and restored after standardization. The workflow then executes an automated sequence of normalization and verification steps designed to correct structural inconsistencies. These include removal of duplicate spatial points, correction of overlapping nodes, pruning of included side branches, and repair of non-positive radius values. Intermediate diagnostic codes are generated and logged to identify the types of irregularities encountered during processing. Standardized files, processing logs, and summaries are written to dedicated output directories. These outputs preserve both the corrected reconstruction and detailed records of the structural issues identified and resolved during the standardization procedure, ensuring reproducibility and traceability of the normalization process.

The second pipeline addresses long connections and disconnected subtrees, which are common artifacts in digital neural reconstructions. After standardization, each SWC file is converted into a directed graph representation using NetworkX [27], where nodes correspond to morphological points and edges represent parent–child connectivity. For each parent–child pair, the Euclidean distance between nodes is calculated. The distribution of connection lengths within a reconstruction is then summarized using mean and standard deviation. Edges whose lengths exceed a user-defined multiple of the standard deviation (6 by default) are considered anomalously long connections. When such edges are detected, the corresponding parent–child relationships are removed and the connectivity of the graph is updated. Following removal of anomalous edges, the pipeline identifies the soma-containing main tree and any disconnected components. Detached subtrees are evaluated relative to the soma, and candidate reconnection points are identified based on spatial proximity and graph topology. If a detached component lies within an adaptive threshold distance, it is automatically reconnected to the main arbor. The process iterates until no disconnected tree remains or a predefined iteration limit is reached. For each reconstruction, the system records statistics including the mean connection length, standard deviation, connection threshold, number of long edges detected, number of edges removed, and number of nodes or subtrees reconnected. These metrics are stored in summaries and control files, enabling detailed auditing of the correction process.

The third pipeline performs automated assignment of pyramidal dendritic labels, distinguishing apical and basal trees within reconstructed neurons. Each SWC file is parsed into a directed morphology graph in which nodes store spatial coordinates, radius values, and parent–child relationships. The soma is identified as the root node, and the arbor is decomposed into subtrees originating from the soma’s direct children. For each candidate subtree, a set of morphology descriptors is computed to characterize its structural and spatial properties. These descriptors include node count, number of bifurcations, maximum Euclidean distance from the soma, maximum root-to-tip path length, total arbor length, Sholl-derived radial complexity, and principal spatial axis orientation. Using these descriptors, candidate dendritic trees are evaluated to determine the most likely apical dendrite, while the remaining eligible trees are classified as basal. Small unbranched stubs and non-dendritic components (e.g., axon) are excluded from the apical candidate set. In addition to the rule-based morphology evaluation, a neural classifier implemented in PyTorch [28] was used to learn tree-level dendritic identity from the extracted morphology features. The model consists of a compact multilayer neural architecture that accepts a feature vector representing each tree and produces a three-class prediction corresponding to apical, basal, or other. For model development, a dataset of 20,500 pyramidal neurons from NeuroMorpho.Org was randomly divided into training, validation, and testing subsets (80/10/10). Training was performed over multiple cycles using the Adam optimizer [29] with adaptive learning-rate scheduling based on validation loss. To accelerate processing, computation was distributed across multiple processes using Message Passing Interface (MPI), allowing parallel parsing of SWC files, feature extraction, and model evaluation. Model performance was assessed using file-level accuracy together with weighted precision, recall, F1-score, and cross-entropy loss.

To support visual verification of neural reconstruction corrections, the pipeline was augmented with an automated PNG rendering workflow, which converted the processed SWC files into images stored alongside the associated log reports.

The entire system, including the web interface, backend processing modules, morphology correction algorithms, neural classification model, and visualization tools, was containerized using Docker and deployed on AWS infrastructure. Centralized logging captured all processing stages, including standardization, structural correction, and dendritic relabeling, while the web interface provided real-time feedback on processing status. This architecture enabled scalable, automated, and reproducible quality control of large neuronal morphology datasets.

## 3. Results

We developed an end-to-end automated quality-control, correction, and standardization framework for SWC-formatted neural morphology reconstructions (Fig. 1). The platform exposes a web-based API that initiates a fully containerized processing pipeline, integrating deterministic structural proofreading, automated normalization, topology repair, geometric correction, morphometric analysis, and machine-learning–based dendritic relabeling within a unified cloud-deployed computational environment.

**Figure 1:**
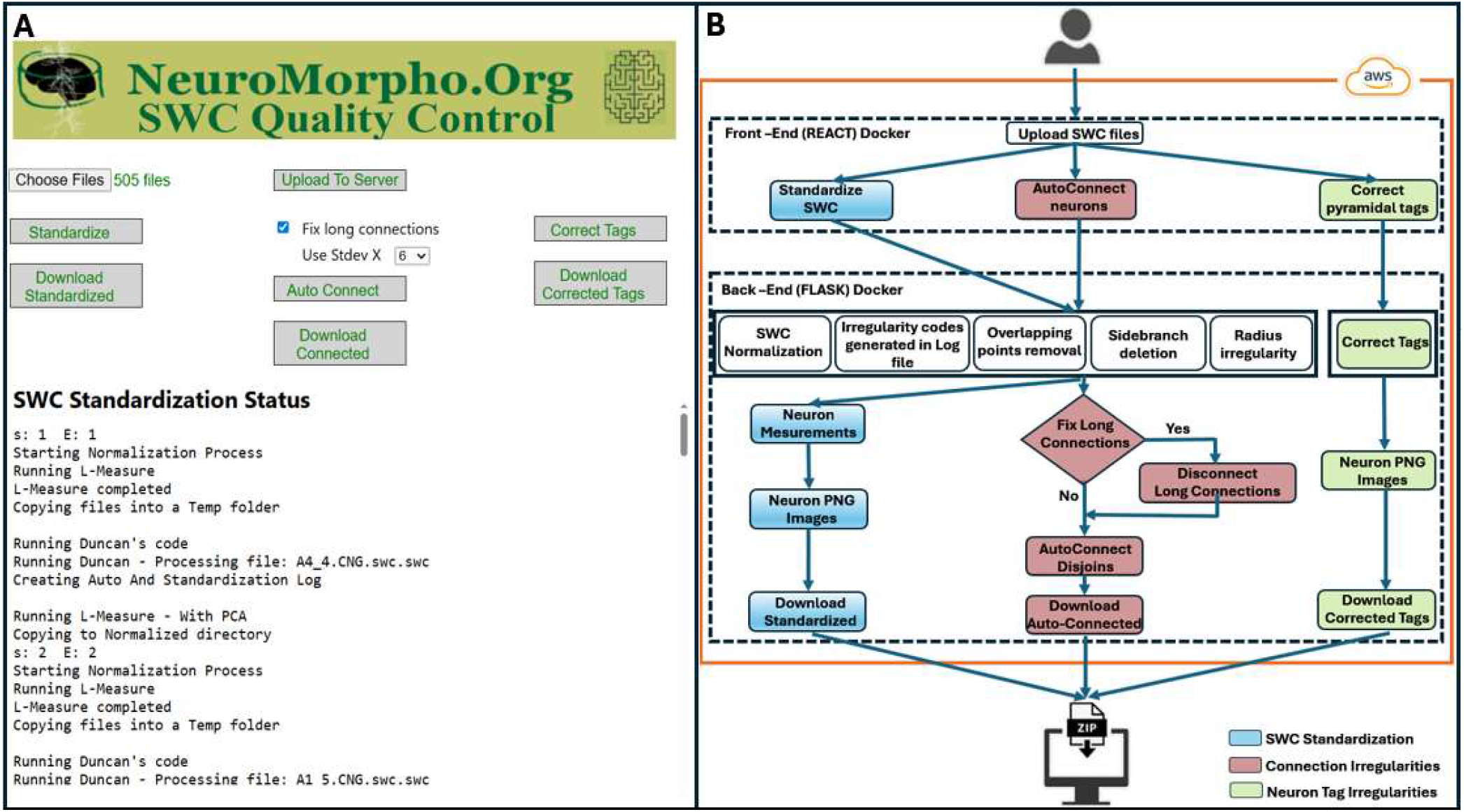
Workflow of the Automated SWC Quality Control and Correction Framework. **A -** Web-based user interface for quality control and standardization of digitally reconstructed neural data. **B -** Workflow of SWC quality control consisting of a containerized React front-end and a Flask back-end interconnected within a Docker network and hosted on AWS Cloud.

Users interact with a React-based front-end to upload SWC files (Fig. 1A). The files are routed through a Flask-based back-end that orchestrates the standardization workflows within isolated Docker containers deployed on AWS (Fig. 1B). In addition to native SWC inputs, other morphology file formats can be incorporated following conversion through xyz2swc [7], which is also accessible via API, enabling compatibility with heterogeneous reconstruction platforms. Upon ingestion, SWC files undergo automated structural validation and formatting using established computational neuroanatomy tools, including L-Measure [5], StdSwc [6], and scripts developed as part of the NeuroMorpho.Org ingestion process [9]. These programs detect geometric and topological irregularities prior to normalization. Identified anomalies, including overlapping nodes, duplicate points, non-positive radii, spurious side branches, disconnected components, and anomalous parent–child relationships, are encoded using conventional irregularity codes [6] and logged for traceability. The correction routines are then executed programmatically, eliminating the need for manual intervention. Long-connection anomalies are identified through Euclidean distance computation within the neural graph representation. Connections exceeding statistically defined thresholds are removed, producing isolated subtrees that are subsequently evaluated and iteratively reconnected based on spatial proximity constraints, ensuring global structural coherence. Following this automated topology repair, standardized reconstructions undergo quantitative morphometric analysis. Compartment-specific measurements are exported as structured datasets, and corrected SWC files are rendered as PNG visualizations for rapid inspection. Final outputs, including standardized morphologies, correction logs, morphometric tables, and visualization files, are packaged for user download. Strict directory isolation, centralized logging, and containerized execution ensure reproducible, transparent, and fully automated quality control of large-scale neuronal morphology datasets while formalizing expert curation logic.

As shown in Fig. 1, the automation framework comprises multiple processing pipelines that share a common standardization core but differ in scope and downstream outputs. The primary (blue) pipeline, after correction of overlapping points, side branches, and radius inconsistencies, computes quantitative measurements and generates PNG renderings, thus serving as the main workflow for ingestion of new data into NeuroMorpho.Org. In contrast, the secondary (red) pipeline performs the same standardization steps while also detecting and removing long-connection artifacts; however, this pipeline does not generate morphometric measurements or visualization outputs. The tertiary pipeline is restricted to dendritic labeling and PNG generation. Therefore, morphometric measurements are produced exclusively within the primary pipeline.

Overlapping points, flagged with code 2.6 in the Standardization Log, arise when two or more nodes occupy identical spatial coordinates, typically due to tracing artifacts or redundant sampling during reconstruction (Fig. 2). Although subtle, such overlaps can distort parent–child relationships and compromise morphometric analyses and electrophysiological simulations. These anomalies are extremely challenging to identify and remove by manual inspection and editing. By systematically detecting and fixing overlapping points at the outset, the framework ensures the integrity and correctness of downstream applications.

**Figure 2:**
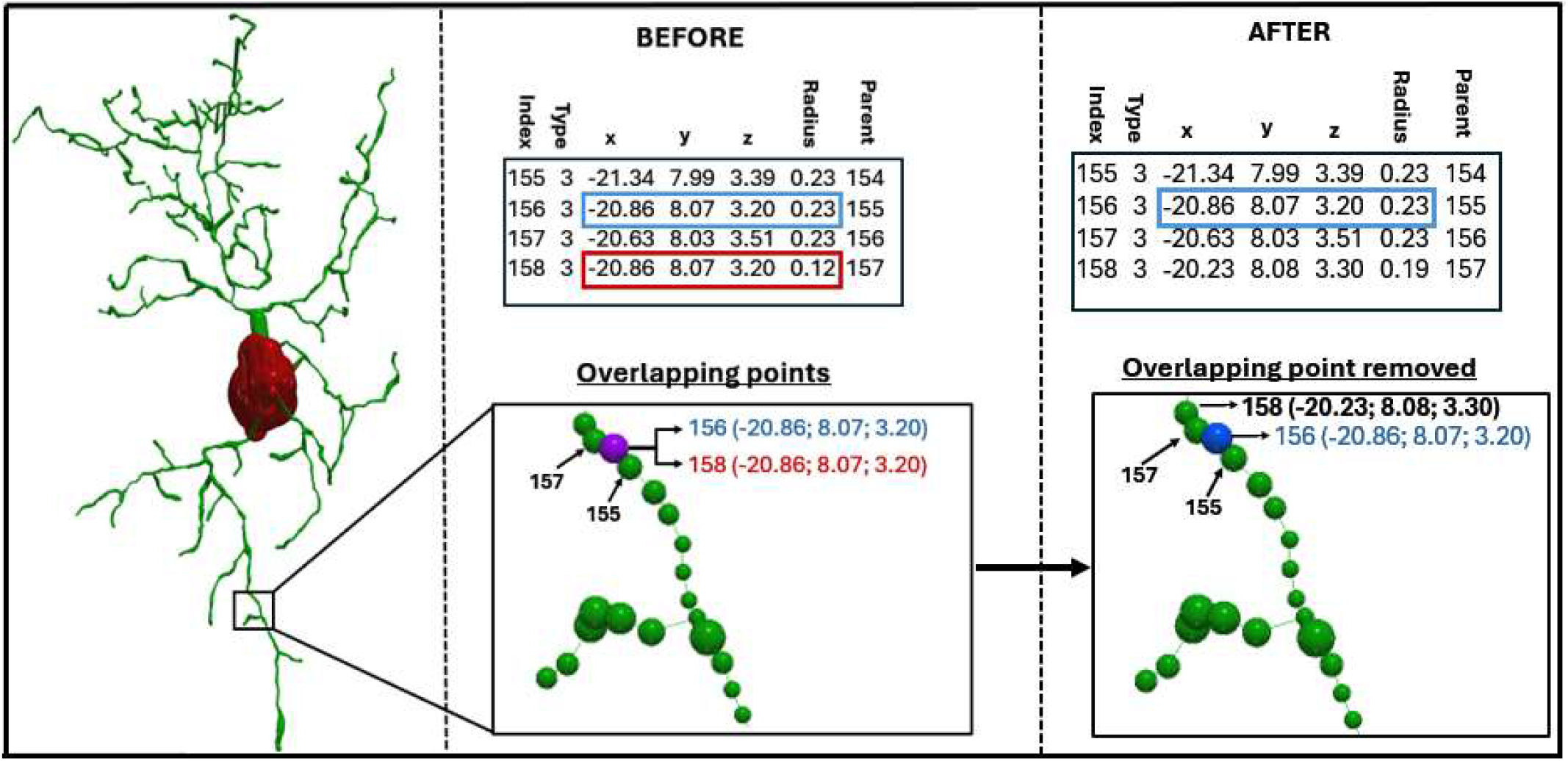
Automated Detection and Correction of Overlapping Points in Neural Morphology Reconstructions. The left panel shows a neuron containing overlapping points detected during the standardization process. The middle panel illustrates two non-adjacent points occupying the same space: blue (index 156) and red (index 158), corresponding to the purple node. The right panel shows the red overlapping point removed after normalization.

Spurious side branches, flagged in the Standardization Log with code 2.7, typically arise when tracing software inadvertently embeds a short collateral stub fully encased within the parent dendritic shaft, creating artifactual skeleton bifurcations that do not reflect true neuronal anatomy (Fig. 3). Such irregularities can distort branching statistics by inflating bifurcation counts and mislead downstream classification models. These artifacts are also difficult to detect visually, but the automated framework resolves them seamlessly through internal graph-based pruning routines without operator intervention.

**Figure 3:**
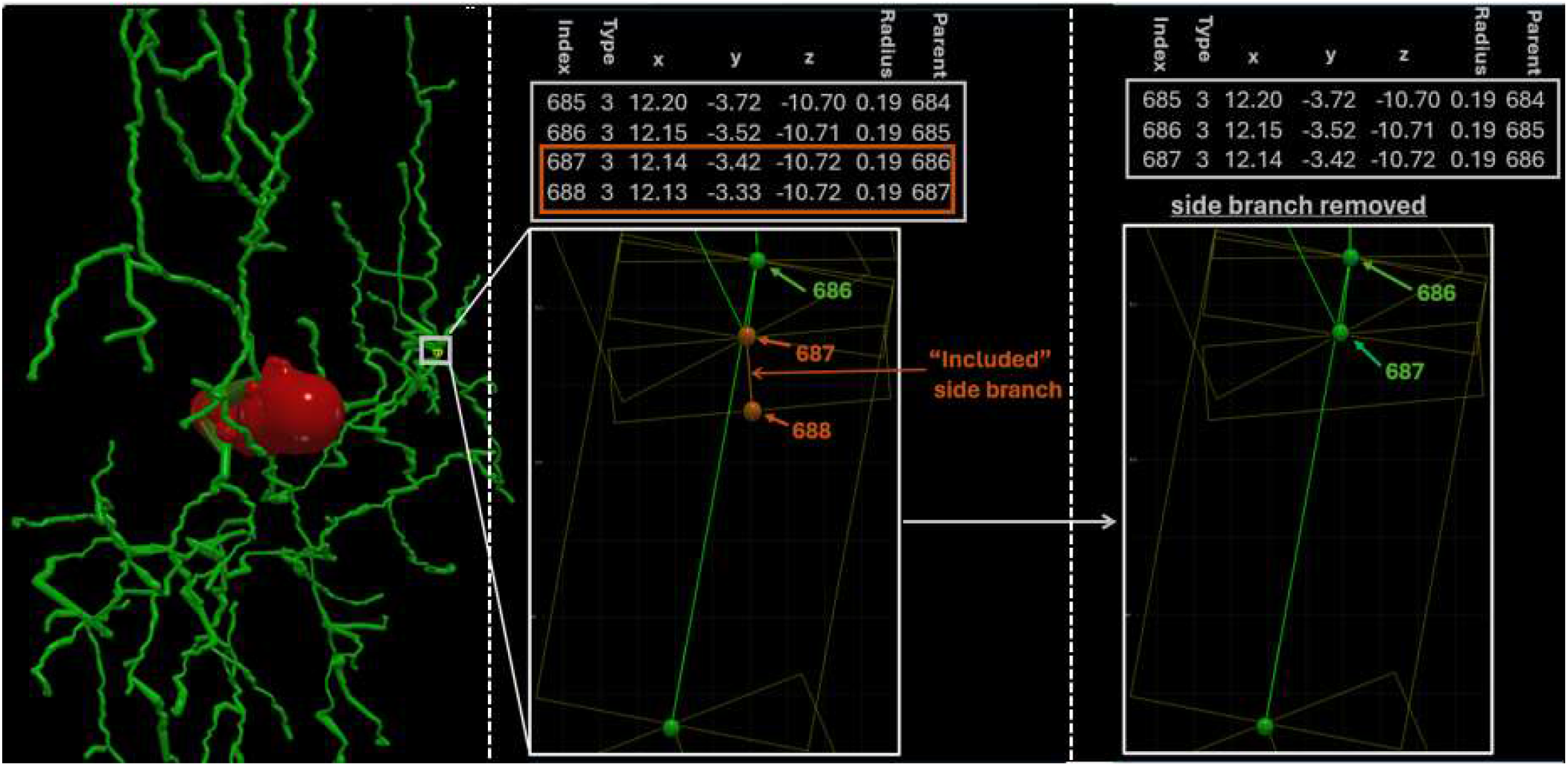
Automated detection and removal of included side branches during SWC morphology normalization. The left panel depicts an included (spurious) side branch detected during the standardization process. The middle panel outlines the small side branch that is not long enough to emerge from within the volume of its parent segment. The right panel shows the side branch removed after normalization.

Non-positive radii, flagged in the Standardization Log with code 4.1, occur when a node is assigned zero or negative thickness during reconstruction, an obvious physical impossibility often produced by data export inconsistencies (Fig. 4). Although seemingly minor, such anomalies can invalidate surface area and occupied volume measurements, in addition to compromising computational simulations. Previously, correcting these inconsistencies required manually executing external utilities such as *stdswc*.*java* [6], introducing additional steps and computing requirements. The automated framework programmatically changes these values to match the radius of the parent node, ensuring biologically valid thickness for all segments prior to downstream analysis.

**Figure 4:**
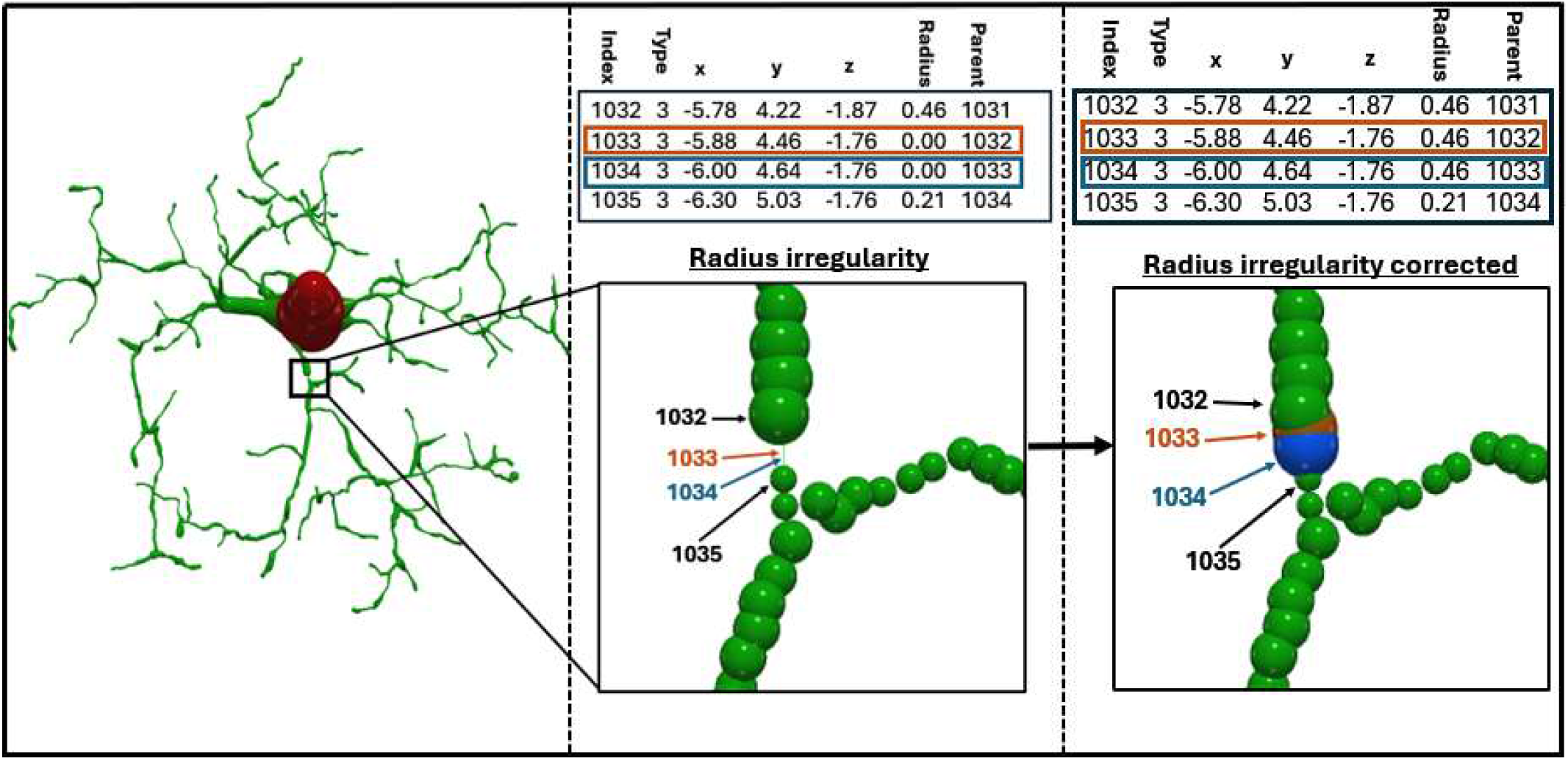
Radius Irregularity Detection and Automated Normalization in Neural Morphologies. The left panel depicts a neuron with radius irregularity detected during standardization. The middle panel illustrates a zero radius at line 1033 shown in brown and line 1034 in blue for display purposes. The right panel portrays the corrected irregularity after normalization with the radius set to that of the parent node.

The first pipeline (SWC Standardization in Fig. 1B) integrates these corrections into a unified framework. Side branches (11% of processed archives) and overlapping points (10% of processed archives) represent two of the most frequently encountered irregularities, consistently ranking among the top five anomaly categories across tracing systems [9]. In the previous NeuroMorpho.Org processing workflow, files were processed sequentially— standardization, overlapping-point removal, spurious side-branch pruning, morphometric computation, and PNG generation—often requiring manual transfers between separate folders at each stage [30]. This fragmented process increased processing time and user interaction (average processing time 62.5 ± 53 min, n=45), introducing labor cost inefficiency and risks of operator error. Consolidating these steps into a single automated pipeline substantially reduces human intervention and eliminates file-handling overhead, decreasing user interaction time at least 30 times while ensuring consistent, reproducible processing across large datasets.

Long connections represent one of the most structurally disruptive reconstruction artifacts. These errors occur when tracing inaccuracies produce incorrect parent–child relationships between nodes that span implausibly large distances not reflective of natural arbor continuity (Fig. 5). Such edges distort neural geometry, compromising morphometric measurements and downstream classification. To address this issue, the application automatically computes the distribution of edge lengths for every reconstruction based on the Euclidean distances between connected nodes. Connections exceeding a user-defined statistical threshold (with default value of six times the standard deviation of inter-node distances) are deemed erroneous and removed, followed by amputation of isolated nodes that are not part of the soma. Because pruning these anomalies fragments the neural arbor, the pipeline then enacts a stitching procedure that identifies the main tree containing the soma and evaluates isolated subtrees based on spatial proximity. Subtrees located within a biologically plausible distance from the soma, main arbor, or each other are reconnected by updating parent–child relationships. This reconnection process is performed iteratively using adaptive thresholding to ensure that all disconnected components are progressively joined while preserving the intrinsic morphological integrity of the original reconstruction (Fig. 5). Throughout this process, quantitative metrics (mean edge distance, standard deviation, number of long connections detected, and number of isolated nodes) are recorded for each file, ensuring that topology repair is not only automated but also transparent and auditable.

**Figure 5:**
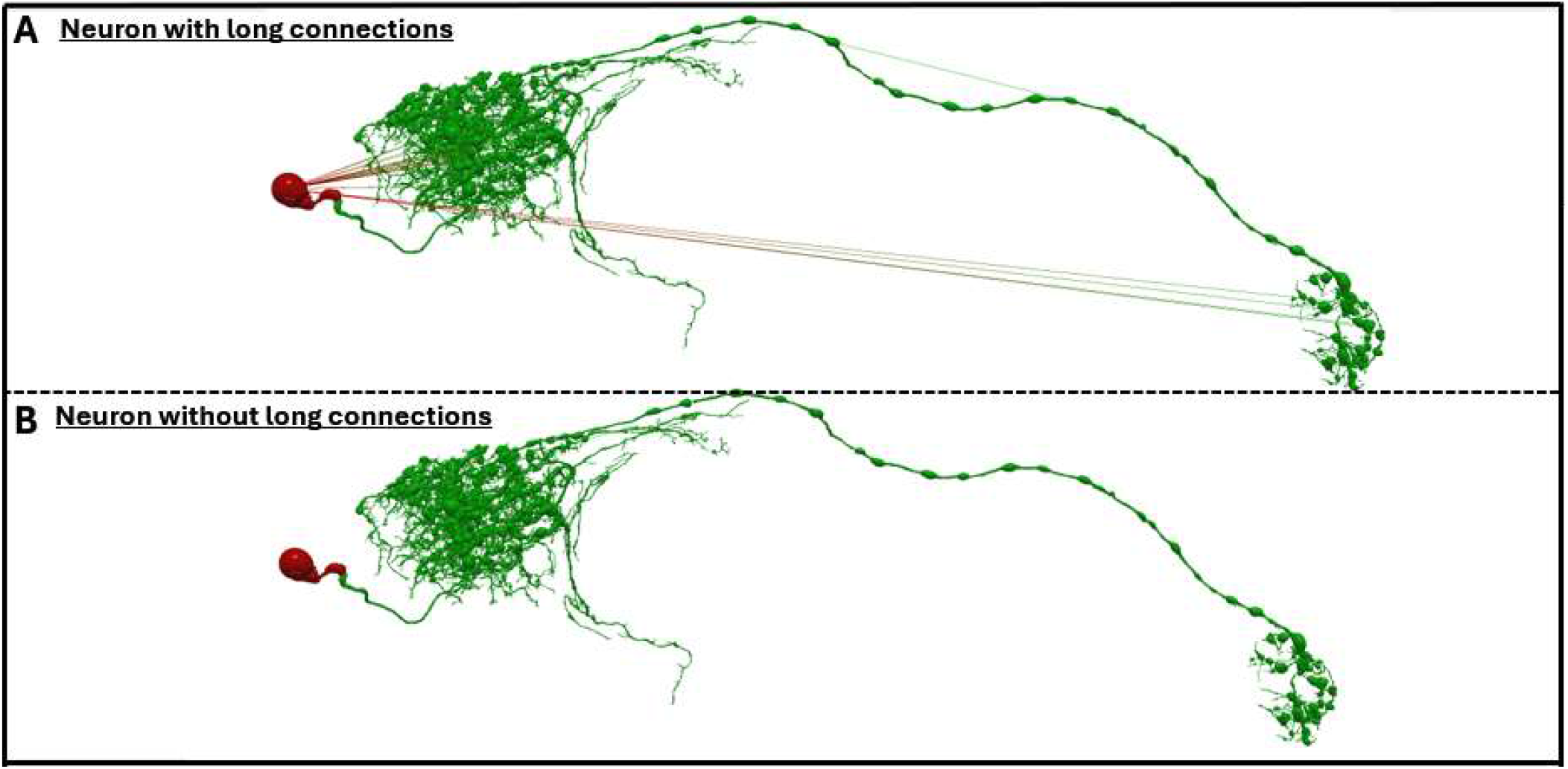
Long Connection Detection and Automated Correction in Neural Morphologies. **A -** Neuron with long connections irregularity having segment lengths greater than the user-defined threshold (here, the default value of 6 times the standard deviation). **B -** From a total of 38,645 connections, 22 connections were considered long connections and removed from the neuron after normalization.

Consistent with prior reports [9], long connections represented the most prevalent structural anomaly, occurring in 29% of processed archives. Before the introduction of this pipeline, correction of these issues required manual editing by a human operator using tools such as Vaa3D [19], NeuTube [21], or TREES [31], typically requiring several minutes for each long connection. In this case, automation has a large impact in terms of efficiency and scalability. In a sample of 264 Flywire reconstructions processed through the long-connection pipeline, an average of 27.1 ± 45.5 long connections per file were corrected in 2.5 ± 25.8 seconds. Importantly, overlapping points were also resolved concurrently (1.9 ± 2.6 per file), meaning that reconstructions emerged from the pipeline fully prepared for morphometric measurement and visualization without additional preprocessing. Tasks that previously required 2 minutes to 1 hour of manual curation per file were thus reduced to seconds through deterministic automation. The framework also enables correction of large-scale reconstructions without compromising morphological fidelity. Manual editing with older computing platforms often requires resampling of today’s larger-sized data file to reduce node density, thereby risking loss of geometric detail. The new automated approach eliminates the need for resampling. For example, four large reconstructions from the Peng archive containing long-connection artifacts were corrected in an average of 39.6 ± 21.4 minutes of computing time, during which 2476.0 ± 606 long connections and 1380.0 ± 414.7 overlapping points were removed per file while preserving the original structural resolution. these cases, the user interaction time was less than 5 minutes for data upload, download, and visual confirmation of accurate processing. Together, these results demonstrate that automated long-connection correction not only accelerates processing but also preserves structural integrity, enabling high-fidelity reconstruction repair at scales previously impractical for manual curation.

Following structural correction and topology restoration, the next critical step is ensuring biologically consistent dendritic labeling. In pyramidal neurons, the presence of a (typically single) apical dendrite stemming from the soma in addition to (generally multiple) basal trees is a defining anatomical feature; however, formatting artifacts and non-specific labeling frequently disregard this distinction. To address this challenge, we applied a graph convolutional network (GCN) trained on 20,500 curated pyramidal neuron reconstructions to identify and correctly tag the apical tree within the overall dendritic arborization (Fig. 6).

**Figure 6:**
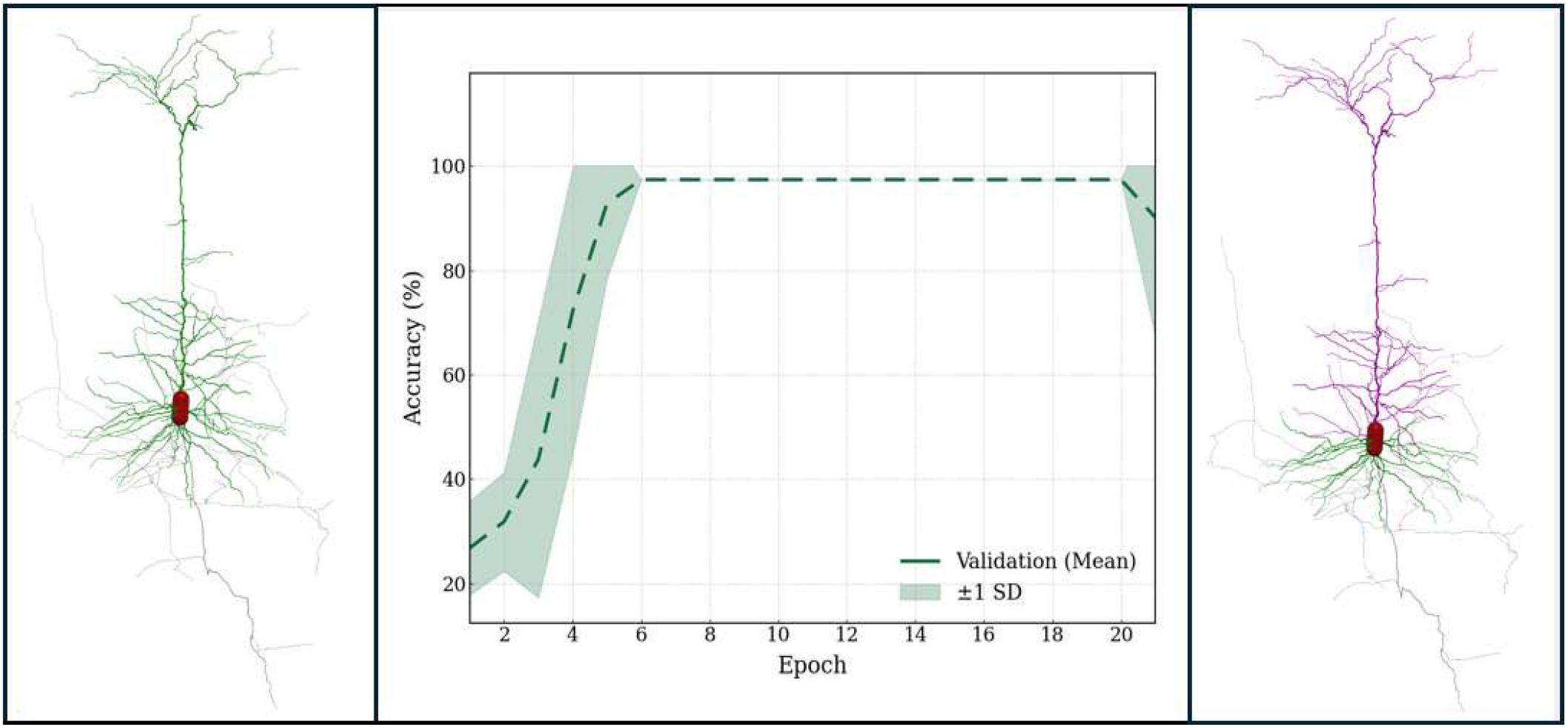
Machine Learning–Based Detection and Automated Correction of Dendritic Tags in Pyramidal Neurons. The left panel shows a pyramidal neuron reconstruction exhibiting uniformly labeled dendrites (green). The middle panel presents the validation accuracy of the machine-learning–based Graph Convolutional Network (GCN) model used for dendrite classification and tagging. The right panel shows the same reconstruction after correction, with basal dendrites (type 3) shown in green, apical dendrites (type 4) in magenta, and the axon (type 2) in gray.

We trained the model over ten independent runs to assess stability and reproducibility. Validation and test accuracies were consistently similar at ∼99.5% in each run, indicating highly stable generalization. At the class level, weighted precision, recall, and F1-scores were 0.978, 0.977, and 0.977, respectively, reflecting balanced performance despite dataset imbalance. Training employed adaptive learning-rate scheduling to ensure stable convergence. To increase biological fidelity, the framework enforced the presence of exactly one apical tree per neuron. Incorporating this constraint improved anatomical consistency without compromising classification accuracy, aligning computational predictions with established cellular morphology. All per-neuron predictions, node sequences, and evaluation metrics are provided as CSV files in the Supplementary Material. All ten training runs were completed in approximately 25 hours, demonstrating scalability to large morphological datasets. Collectively, these findings indicate that the GCN-based relabeling module achieves highly accurate dendritic classification while preserving biologically grounded constraints and maintaining reproducible performance across independent training runs.

## 4. Discussion

The present work demonstrates that automated, end-to-end quality control and dendritic relabeling for neural reconstructions is both feasible and highly reliable at scale. By integrating standardized morphology-processing tools with a deep graph-based model and a cloud-native orchestration framework, we bridge a longstanding gap between manual curation practices and the computational demands of modern neuroinformatics pipelines.

A key strength of this system is the combination of deterministic structural correction algorithms with probabilistic classification for biological consistency. Traditional standardization utilities efficiently identify and repair syntactic or geometric irregularities, yet require substantial manual oversight and fragmented execution environments. In contrast, the current architecture centralizes these steps, tracks each operation, and secures reproducibility through containerization, directory isolation, and automated logging. This integration reduces operator burden and minimizes human variability, addressing a critical bottleneck when managing large or continuously updated neural data collections. The importance of scalable and automated processing is further underscored by the emergence of massive connectomics and neuroanatomical datasets containing millions of reconstructed neuronal segments and synaptic connections, such as those produced by the FlyWire project, the Brain Initiative Cell Census Network, the MICrONS program, and other recent large-scale reconstruction efforts, which collectively highlight the need for reliable automated pipelines capable of handling increasingly complex and high-volume morphology datasets [32-35].

The reliability of the graph convolutional network in identifying biologically plausible dendritic labeling also represents a meaningful advance. Pyramidal neurons, with their characteristic apical dendrite morphologically and biophysically distinct from the basal trees, represent the most abundant excitatory neuronal class in the mammalian cerebral cortex of which they constitute the sole long-range projecting output across multiple species [36-39]. This biological prominence is reflected in NeuroMorpho.Org, where pyramidal neurons account for nearly one-fifth of the entire database content. The distinction between apical and basal trees is functionally critical, as these dendritic domains receive different afferent inputs and support specialized computational roles [40-42]. Basal dendrites primarily integrate local intracortical and feedforward inputs, whereas apical dendrites are preferentially targeted by long-range, feedback, and modulatory projections, contributing to hierarchical processing, contextual modulation, and learning-related plasticity [40,41]. Moreover, basal and apical trees follow specialized developmental cues [22], and the axon can originate from basal, but not apical dendrites [43]. Accurate identification of these compartments is therefore essential for faithful morphometric analysis, biologically grounded circuit modeling, and interpretation of domain-specific synaptic integration. By enabling automated and reliable apical–basal delineation at scale, our framework supports large-scale studies of excitatory neuron structure–function relationships across species and brain regions. Ensuring accurate and automated identification of apical branches enables more faithful downstream computational simulations, morphometric comparisons, and circuit analyses. While the performance outcomes are strong, opportunities remain for continued development. Expanding the feature space beyond Sholl-based metrics may improve generalization to additional structural domains (e.g., axons) and neuron types [44]. Incorporating unsupervised components or self-supervised learning may further enhance robustness in settings with incomplete or noisy training labels.

Importantly, the cloud-oriented design ensures that the system is not limited to a single laboratory or compute setting. The use of Docker and AWS allows consistent deployment across institutions, enabling scalable and collaborative workflows. Such portability paves the way for community-wide implementation in shared data repositories, imaging facilities, and automated reconstruction pipelines, ultimately enhancing transparency, reproducibility, throughput, and fair credit assignment of morphological datasets worldwide [45]. Overall, this platform provides a scalable and reproducible foundation for the next generation of neuron morphology curation. This capacity will be increasingly critical as emerging imaging technologies and connectomics programs continue to accelerate data acquisition far beyond the limits of manual review. The system represents a practical step toward fully automated morphological curation and sets the stage for broader integration of artificial intelligence in structural neuroscience workflows.

## 5. Conclusions

This study introduces a fully automated, cloud-enabled pipeline for the standardization, correction, and biological refinement of neuronal morphology reconstructions. By unifying established morphology-processing tools, reproducible containerized infrastructure, and a high-accuracy graph convolutional network for dendritic relabeling, the system eliminates key manual bottlenecks in SWC data curation and ensures consistent, scalable processing across large datasets. This framework provides a practical foundation for future automated neuroinformatics infrastructures, facilitating rapid, reproducible, and standardized preparation of morphological data for downstream computational analysis, atlas construction, and large-scale neuroscience initiatives.

## Supporting information

Standardization and accuracy results

## Availability of Data and Materials

The digital reconstructions of neural morphology data were fetched from the publicly available repository NeuroMorpho.Org. The source code and datasets for the Standardization and Quality Control of the digital reconstructions of neural morphology are available at https://github.com/HerveEmissah/nmo_swc_standardization. The executable pipeline is accessible at https://swcstandardization.computational-neuromorpho.org.

## Author Contributions

H.E. conceived and designed the study, developed the computational framework and algorithms, implemented the software pipeline, performed the data processing and analysis, and drafted the manuscript.

C.T. contributed to data curation, validation, and interpretation of the results, and assisted in reviewing and editing the manuscript.

G.A.A. conceived and designed the study, provided conceptual guidance, supervision, and critical revisions of the manuscript.

All authors contributed to editorial revisions, read, and approved the final version of the manuscript.

## Ethics Approval and Consent to Participate

Not applicable.

## Acknowledgment

This research was supported by resources provided by the Office of Research Computing at George Mason University (URL: https://orc.gmu.edu) and funded in part by grants from the National Science Foundation (Award Number 2018631).

The authors are grateful to Drs. Duncan Donohue, Sridevi Polavaram, and Sumit Nanda for developing various components of the NeuroMorpho.Org data processing codebase, which were used in this work.

## Declaration of AI and AI-assisted Technologies in the Writing Process

During the preparation of this work the authors used ChatGpt-5.1 in order to check spelling and grammar. After using this tool, the authors reviewed and edited the content as needed and take full responsibility for the content of the publication.

