## Supplementary figures and images for "Automated Proofreading of Digitally Reconstructed Neural Morphology Enhances Accuracy, Scalability, and Standardization"

### 720575940616379707.png

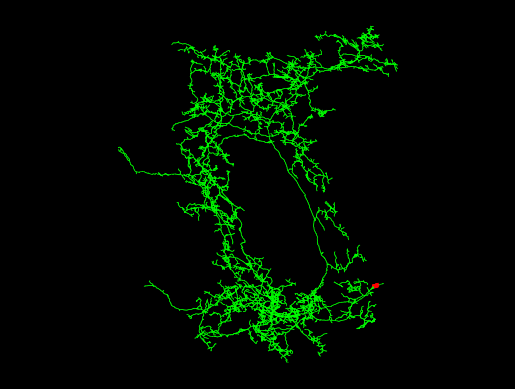

### 720575940616383193.png

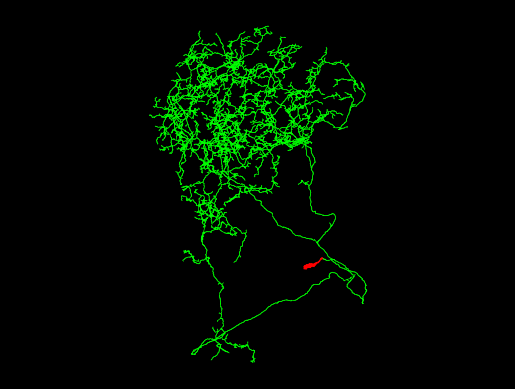

### 720575940616390261.png

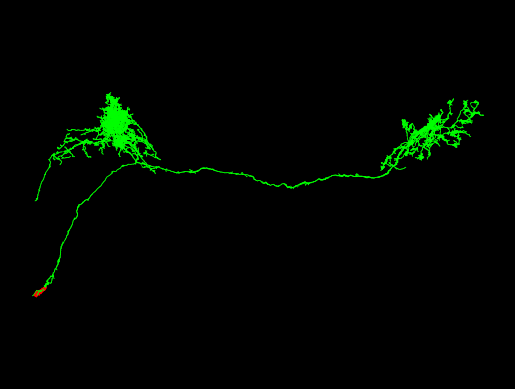

### 720575940616392849.png

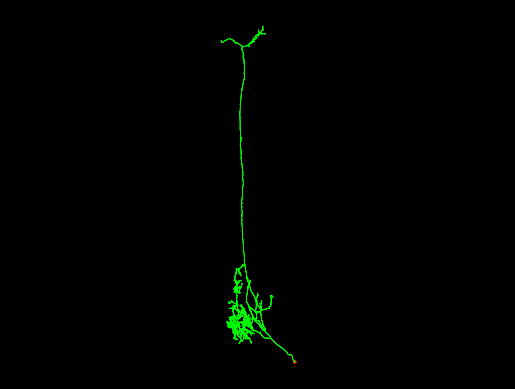

### 720575940616399645.png

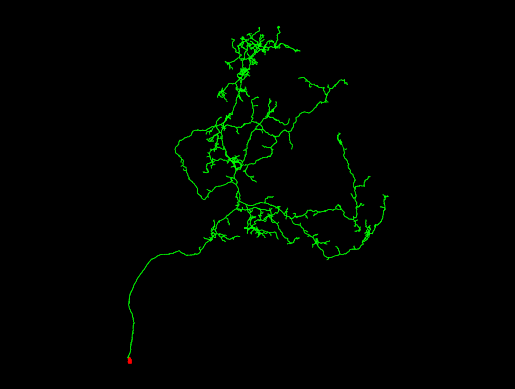

### 720575940616402269.png

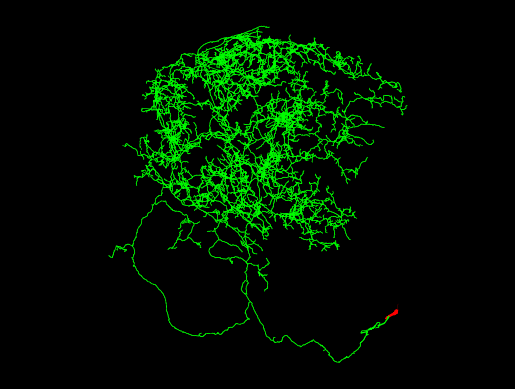

### 720575940616408065.png

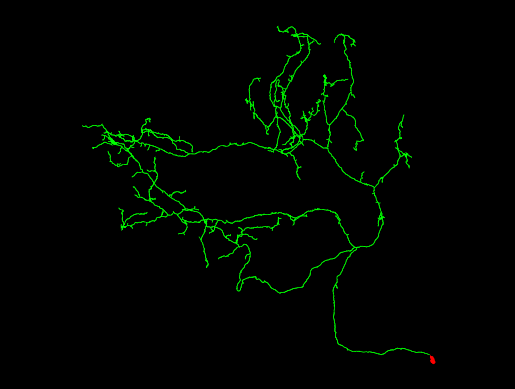

### 720575940616408262.png

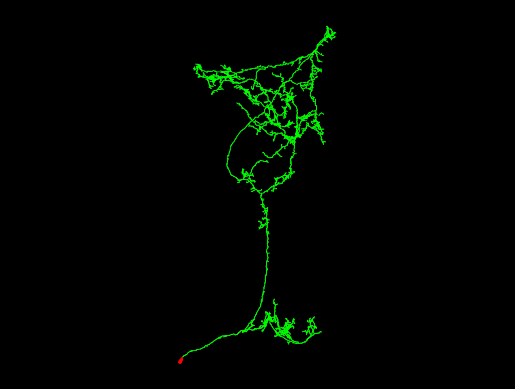

### 720575940616410171.png

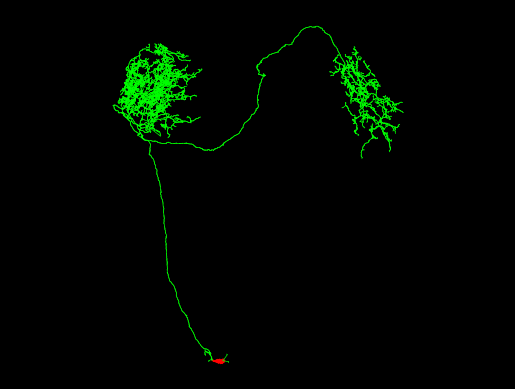

### 720575940616410566.png

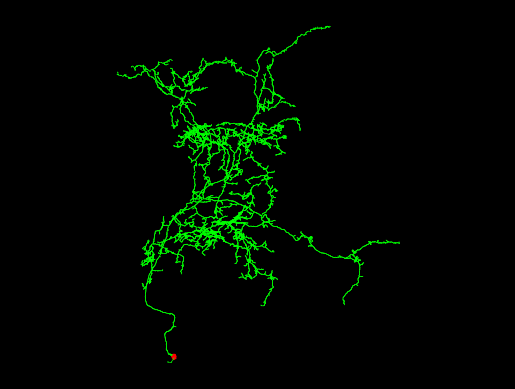

### 720575940616413894.png

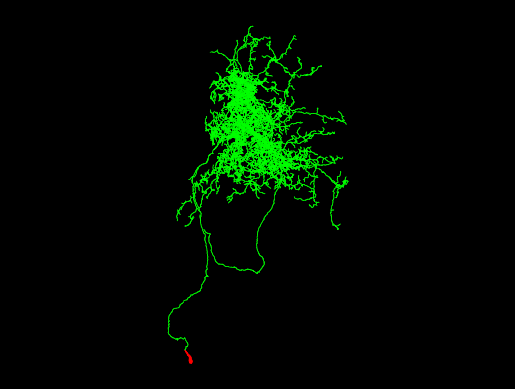

### 720575940616415003.png

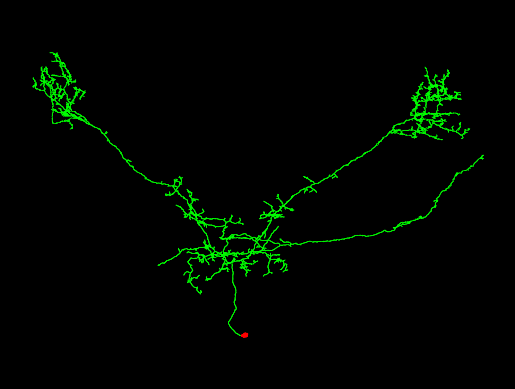

### 720575940616420865.png

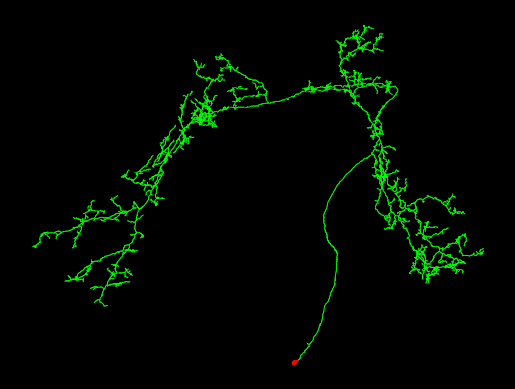

### 720575940616425531.png

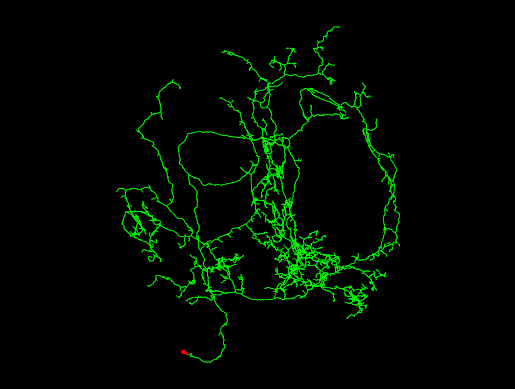

### 720575940616429825.png

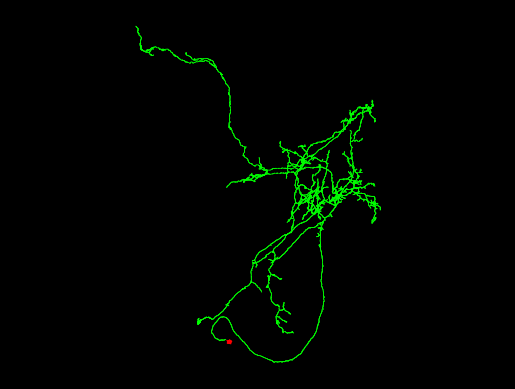

### 720575940616441181.png

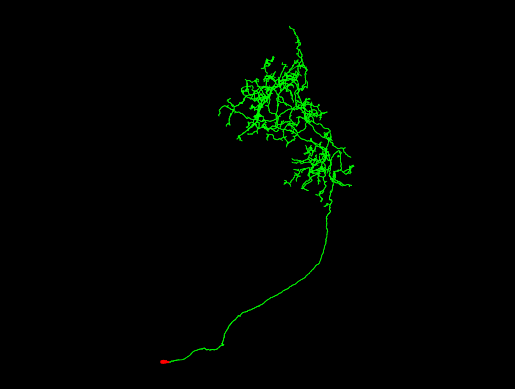

### 720575940616448581.png

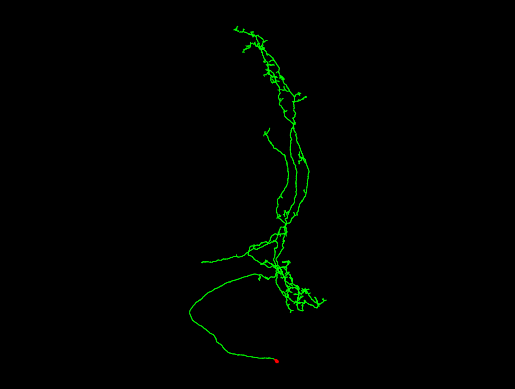

### 720575940616451909.png

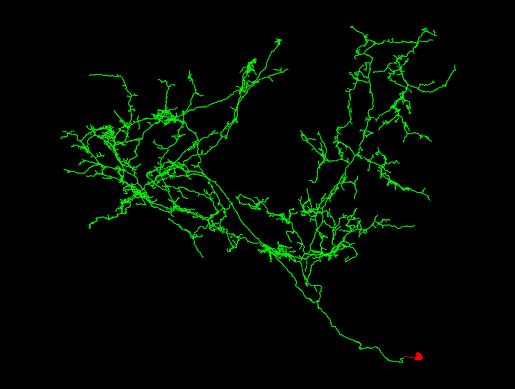

### 720575940616459333.png

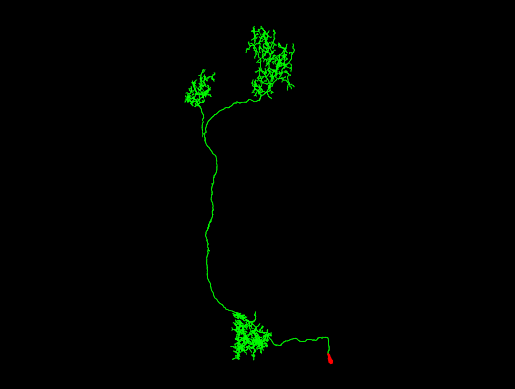

### 720575940616462987.png

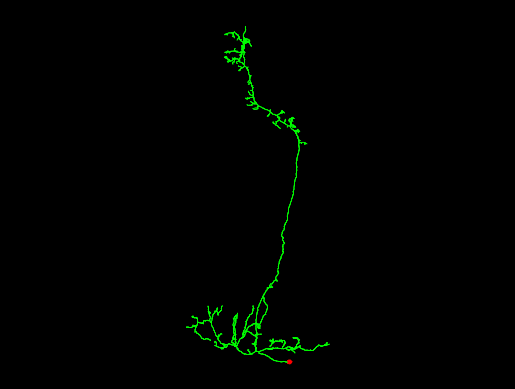

### 720575940616463477.png

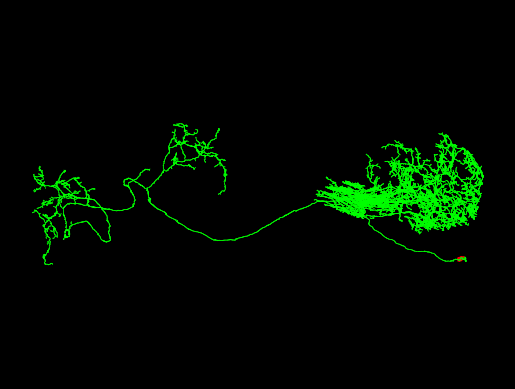

### 720575940616464340.png

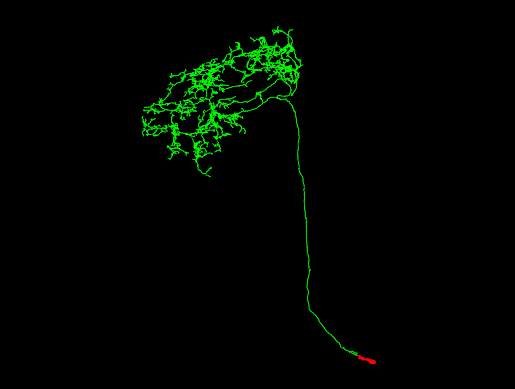

### 720575940616466260.png

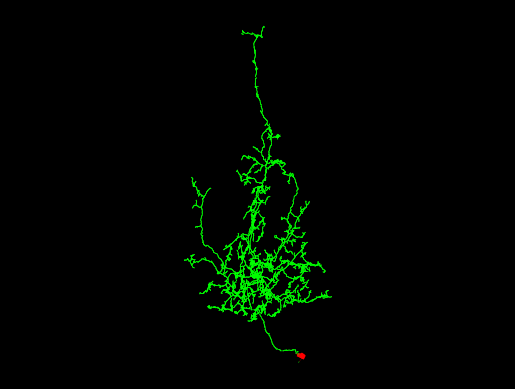

### 720575940616466388.png

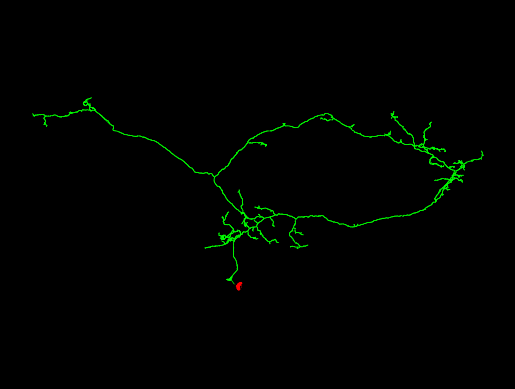

### 720575940616466805.png

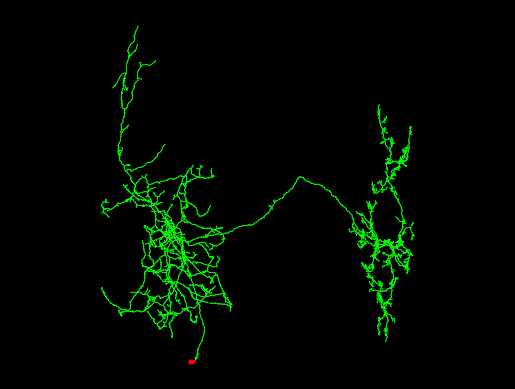

### 720575940616481397.png

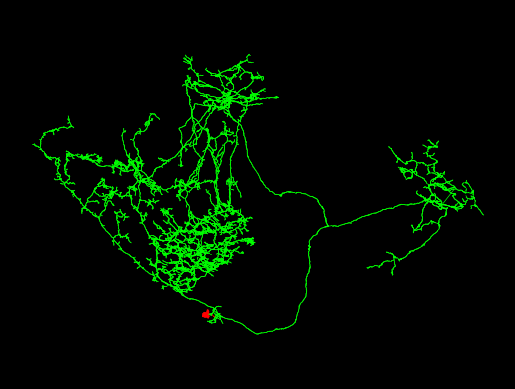

### 720575940616483284.png

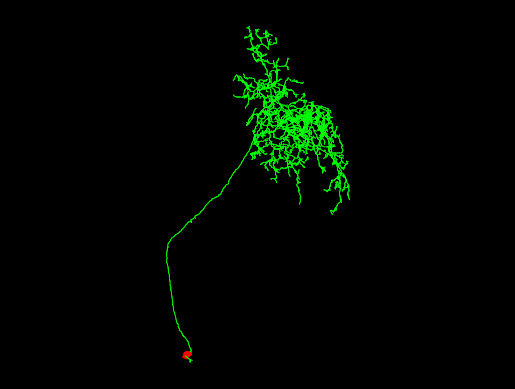

### 720575940616483414.png

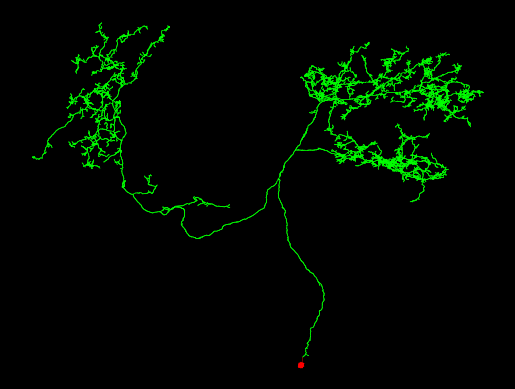

### 720575940616486230.png

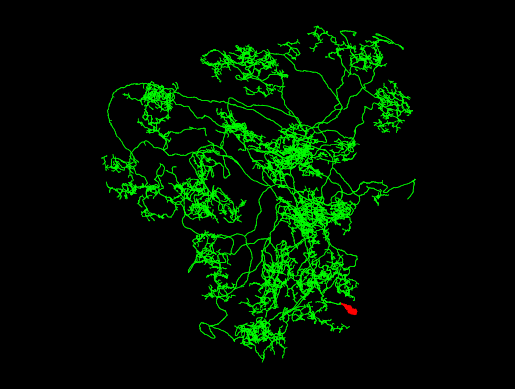

### 720575940616501877.png

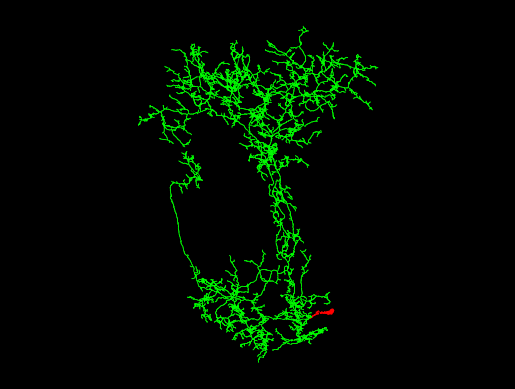

### 720575940616502109.png

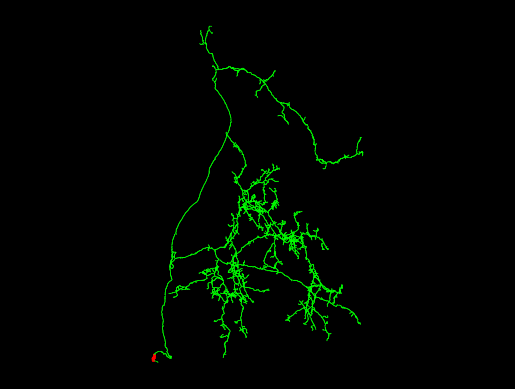

### 720575940616504977.png

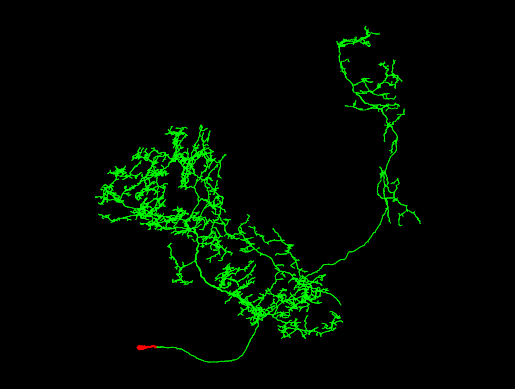

### 720575940616505017.png

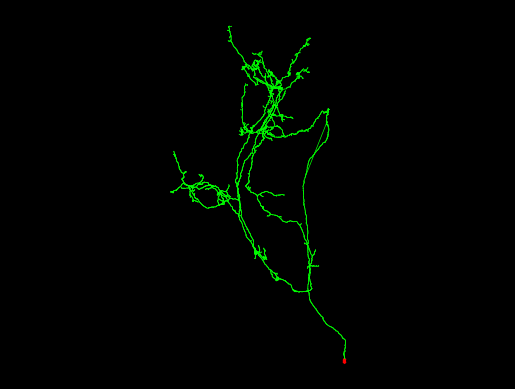

### 720575940616507865.png

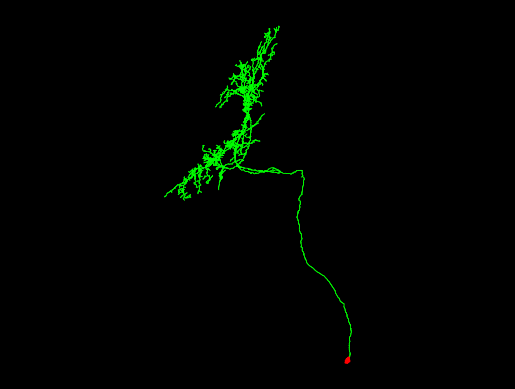

### 720575940616511813.png

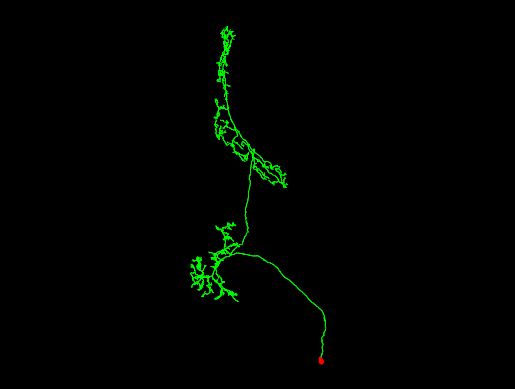

### 720575940616525429.png

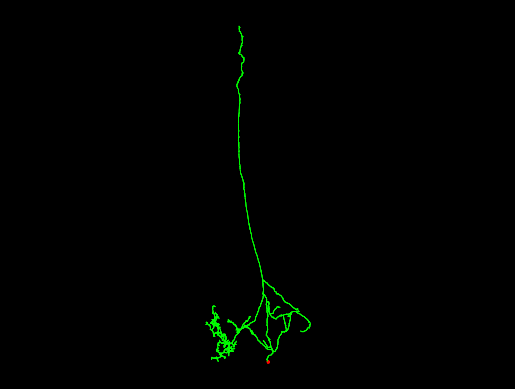

### 720575940616538944.png

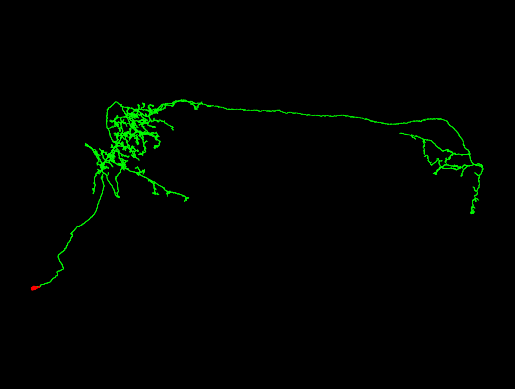

### 720575940616539990.png

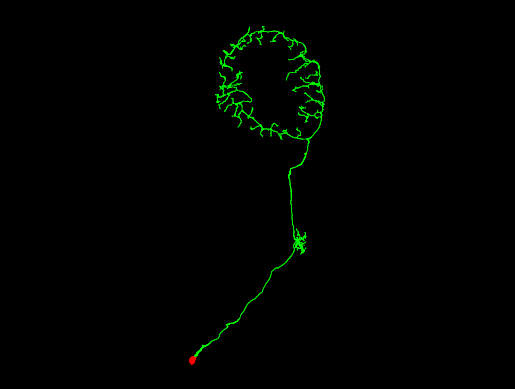

### 720575940616543745.png

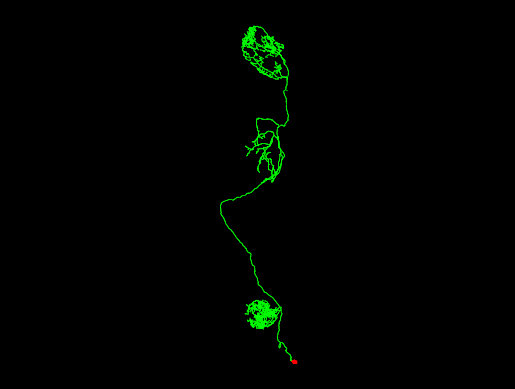

### 720575940616566133.png

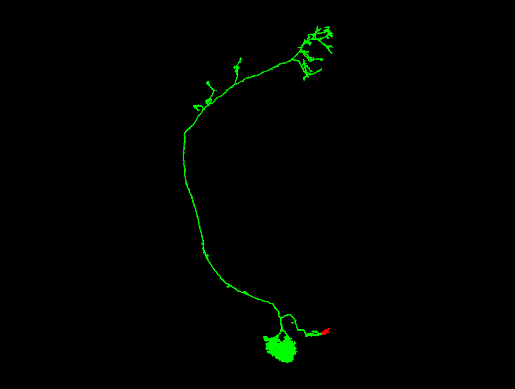

### 720575940616702465.png

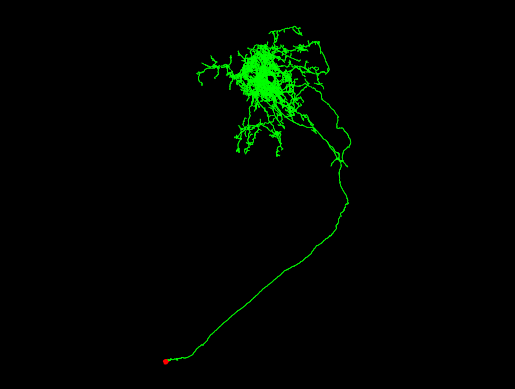

### 720575940616848214.png

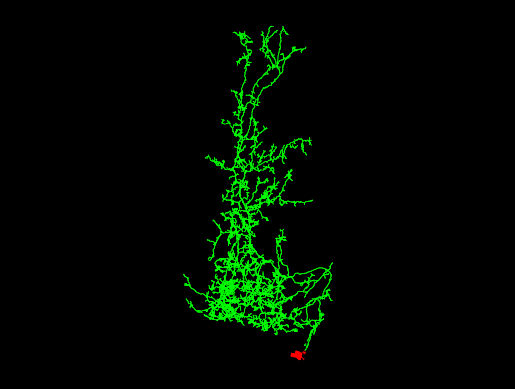

### 720575940617422641.png

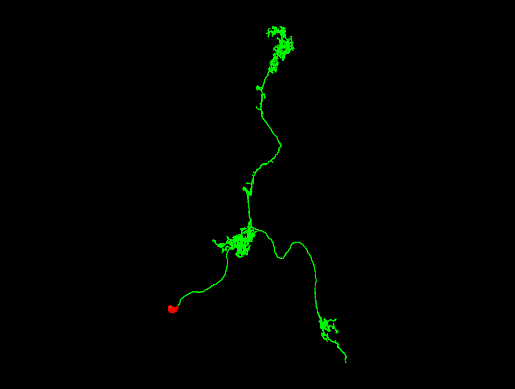

### 720575940617425593.png

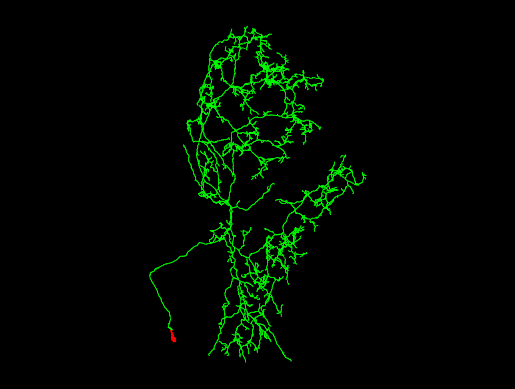

### 720575940617429853.png

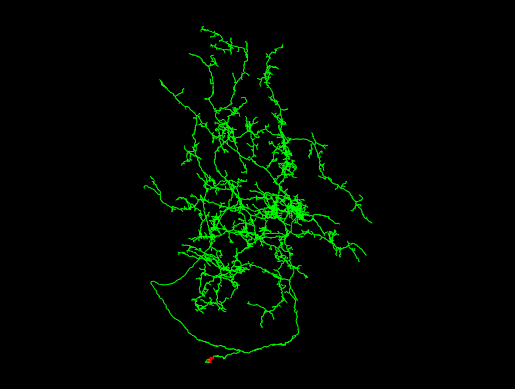

### 720575940617431325.png

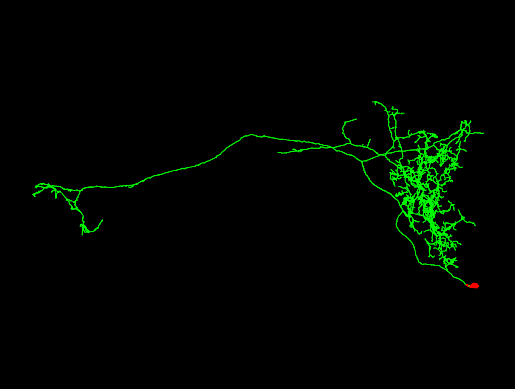

### 720575940617433627.png

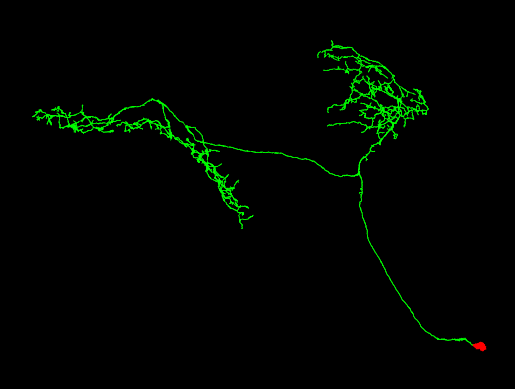

### 720575940617440395.png

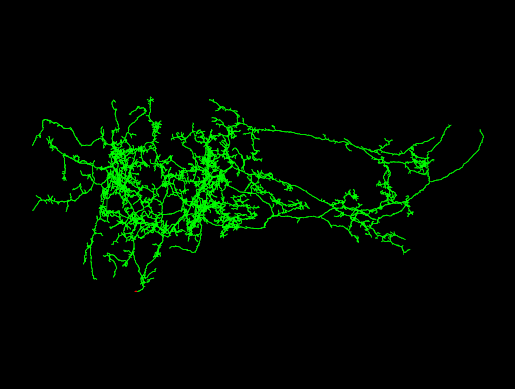

### 720575940617440802.png

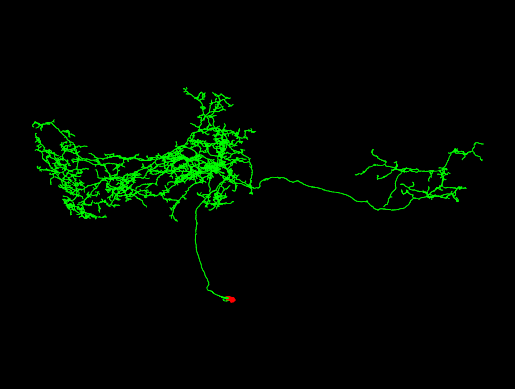

### 720575940617441752.png

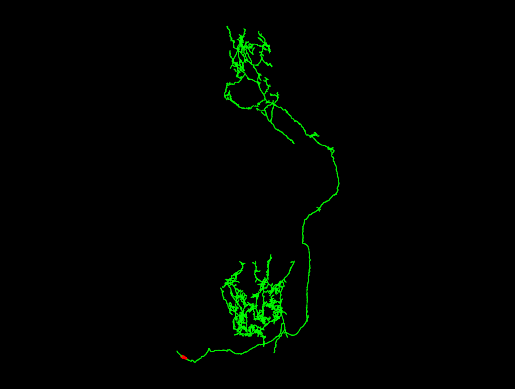

### 720575940617452500.png

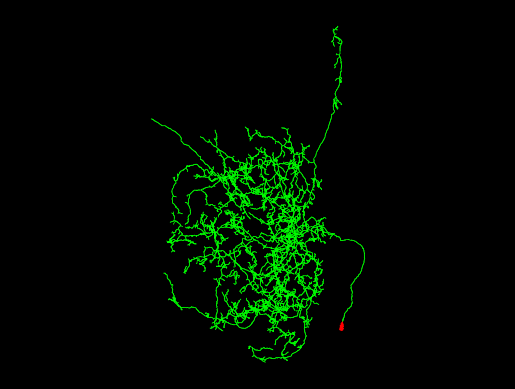

### 720575940617466395.png

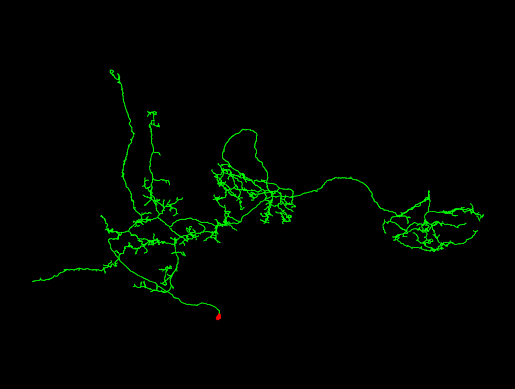

### 720575940617467712.png

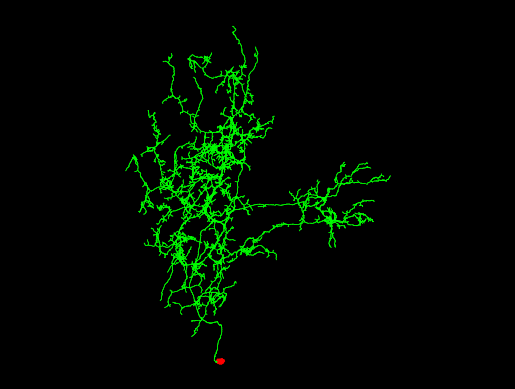

### 720575940617470164.png

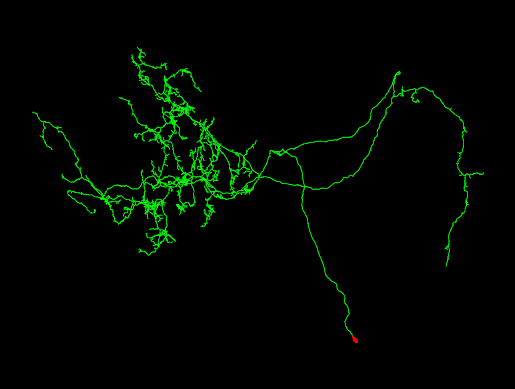

### 720575940617470932.png

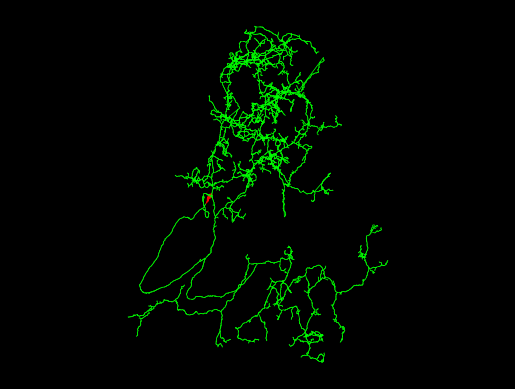

### 720575940617475028.png

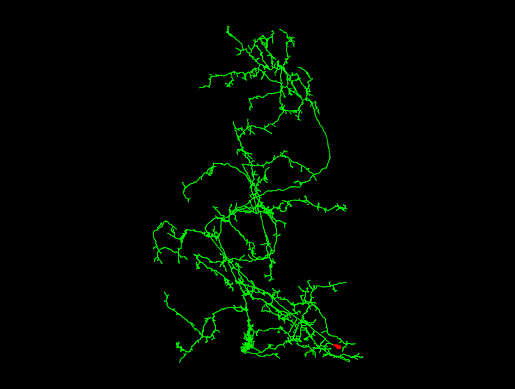

### 720575940617476052.png

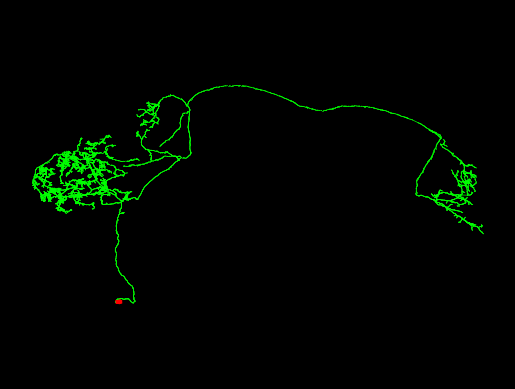

### 720575940617476379.png

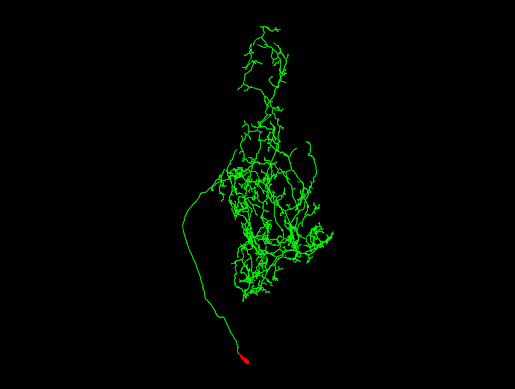

### 720575940617479202.png

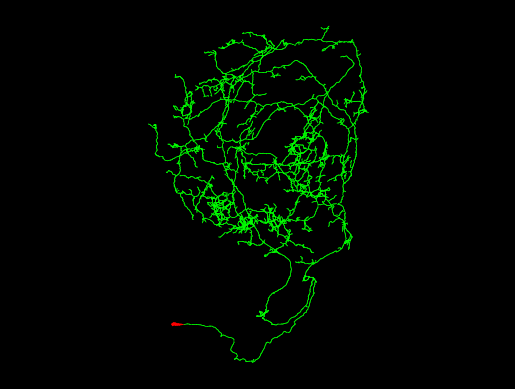

### 720575940617479707.png

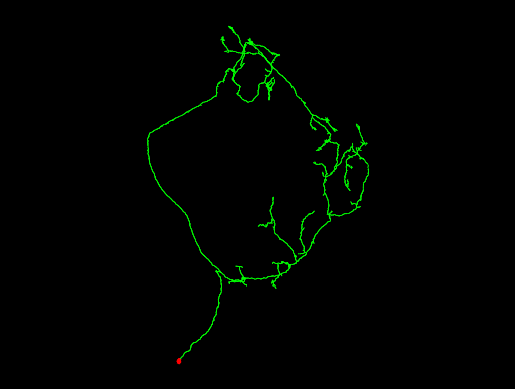

### 720575940617488477.png

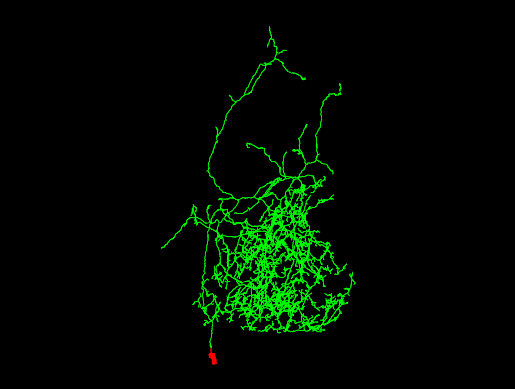

### 720575940617489409.png

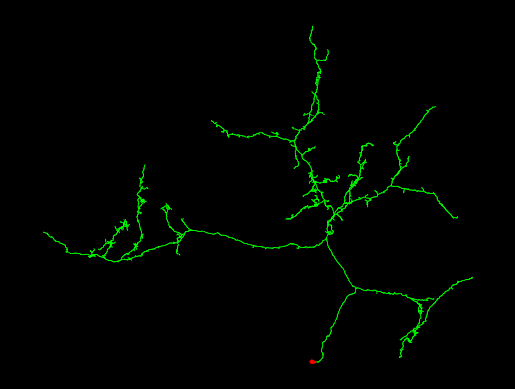

### 720575940617492821.png

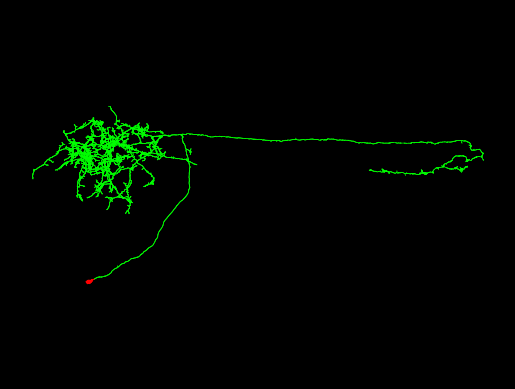

### 720575940617493460.png

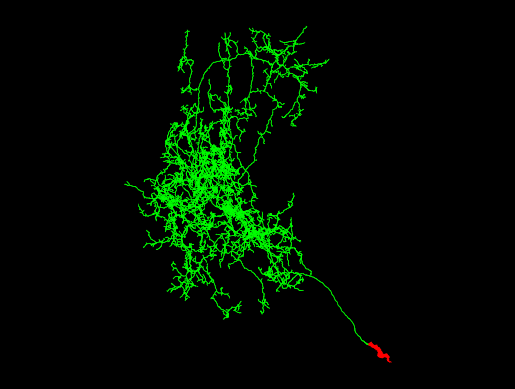

### 720575940617494417.png

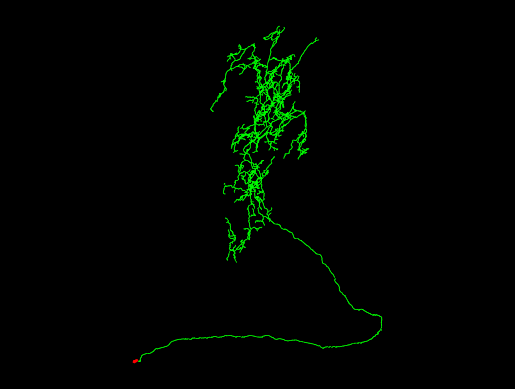

### 720575940617494877.png

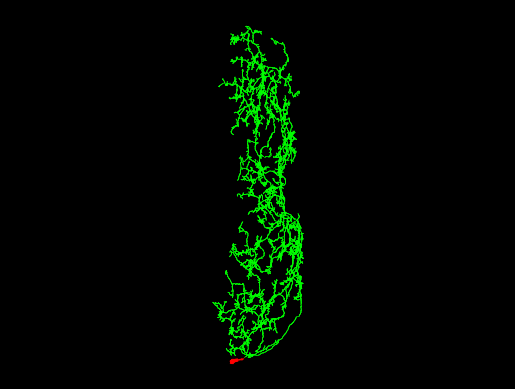

### 720575940617496209.png

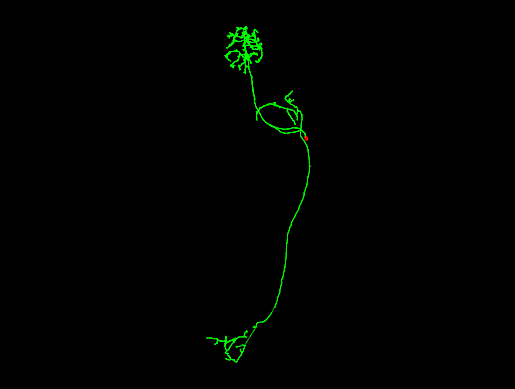

### 720575940617496861.png

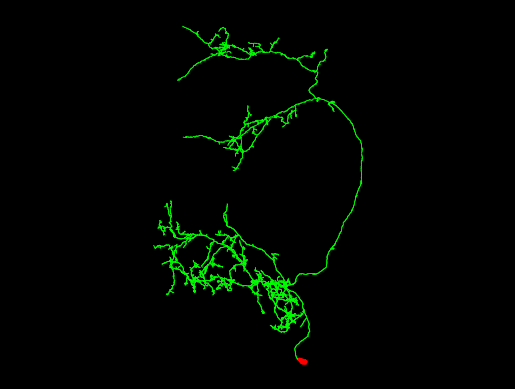

### 720575940617496925.png

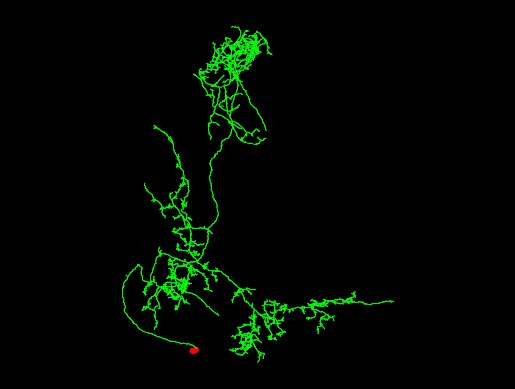

### 720575940617497437.png

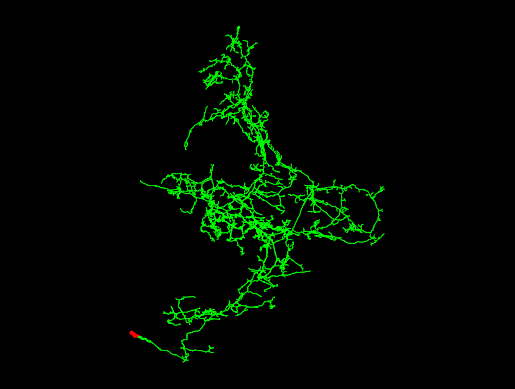

### 720575940617498717.png

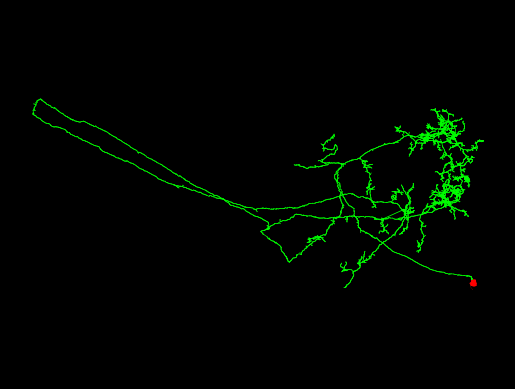

### 720575940617499205.png

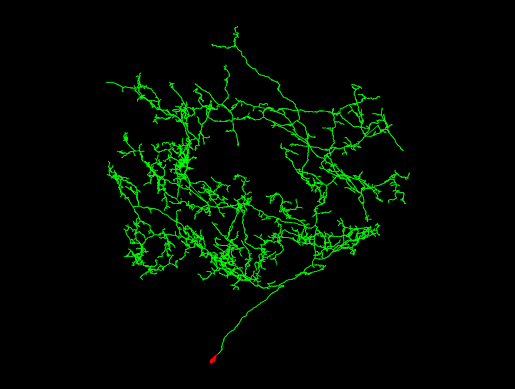

### 720575940617501533.png

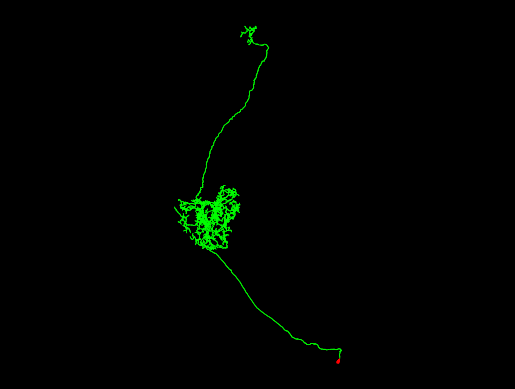

### 720575940617503837.png

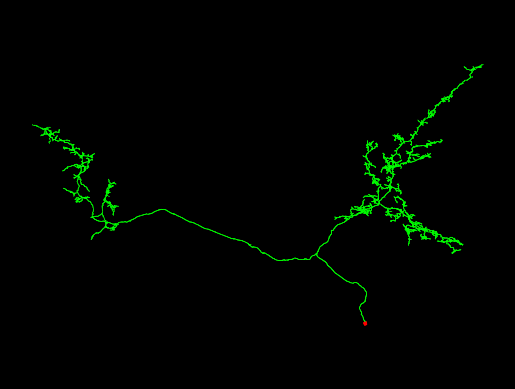

### 720575940618374558.png

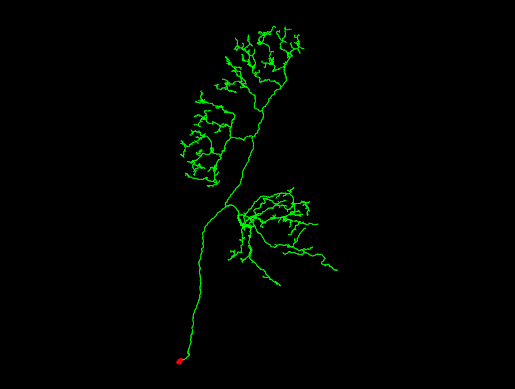

### 720575940618380667.png

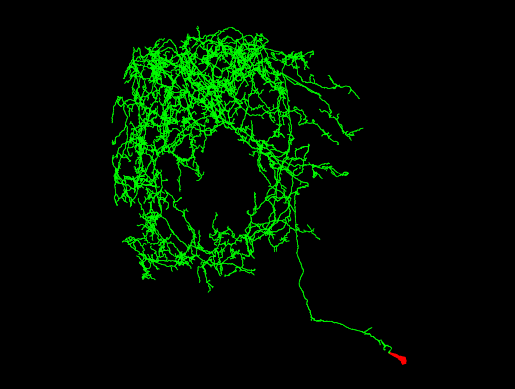

### 720575940618516894.png

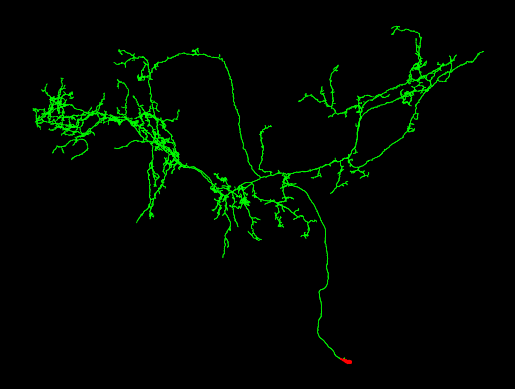

### 720575940618518582.png

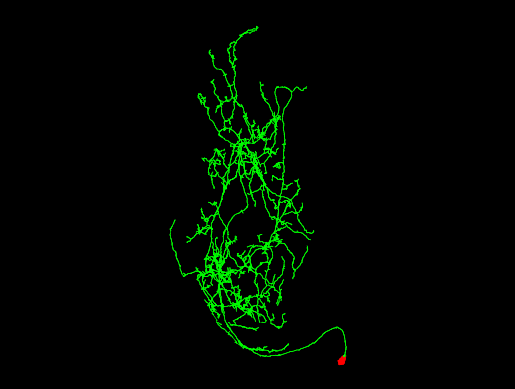

### 720575940618522673.png

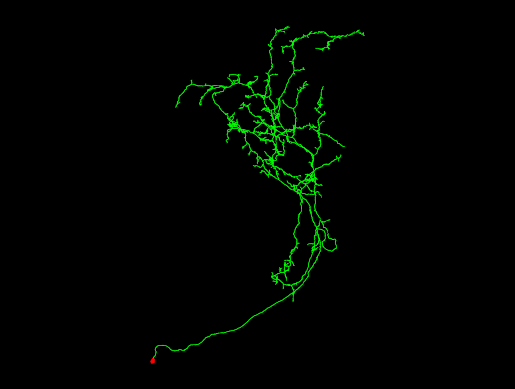

### 720575940618524481.png

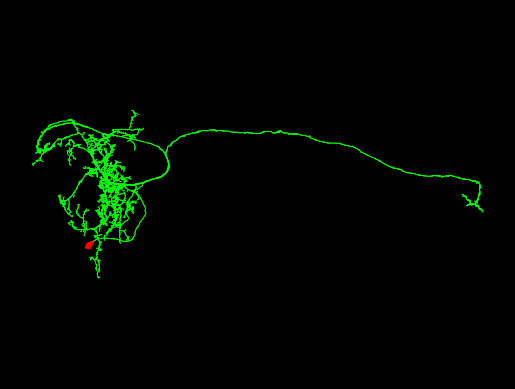

### 720575940618532779.png

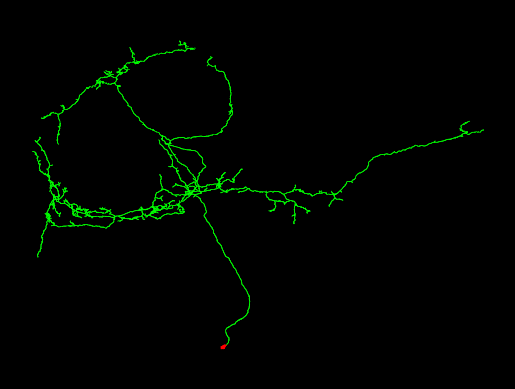

### 720575940618536528.png

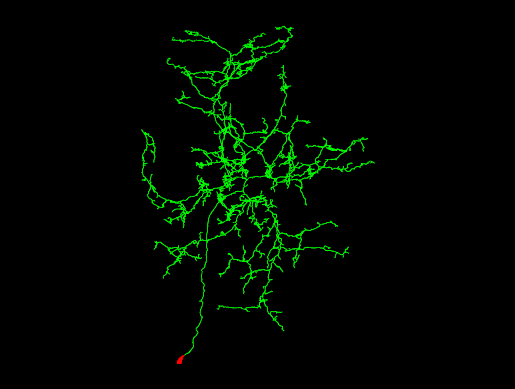
